# Temporal changes in miR-23a-5p levels in subclinical atherosclerosis: A Potential biomarker

**DOI:** 10.1101/2025.07.24.663880

**Authors:** Abinayaa Rajkumar, Gokulprasanth Panchalingam, Deepthy Jayakumar, Devasimman Perumal, Pazhanisankar Muthusamy, Damodharan Shankar, Divya Bhavani Ravi, Kalaiselvi Periandavan

## Abstract

A high-cholesterol diet (HCD) is a well-established contributor to oxidative stress, inflammation, and atherogenesis, yet the temporal progression of these pathological changes and the regulatory role of circulating microRNAs (miRNAs) remain incompletely understood. In this study, we investigated the effects of prolonged HCD exposure to rats on lipid metabolism, oxidative stress, inflammation, endothelial dysfunction, and miRNA expression in a rat model. Male Wistar rats were fed either a standard diet or HCD for 30, 60, 90, or 120 days. Biochemical analyses revealed a progressive dyslipidemic profile characterized by elevated total cholesterol, triglycerides, LDL, VLDL, and atherogenic index, alongside a significant reduction in HDL. Lipid peroxidation levels increased markedly with HCD duration, while antioxidant markers PON1 and ApoA1 were significantly reduced and oxLDL levels were elevated. Inflammatory cytokines (CRP, TNF-α, IL-6, IL-8) and VCAM-1 were significantly increased, indicating systemic inflammation and endothelial activation. Notably, miR-23a-5p and miR-15b-5p were elevated in the serum of HCD-fed rats, with miR-23a-5p peaking at day 120. Correlation analyses revealed that miR-23a-5p expression positively correlated with oxLDL and inversely with ApoA1 and PON1 levels, suggesting its involvement in oxidative and lipoprotein dysregulation. These findings demonstrate that prolonged HCD induces a coordinated cascade of oxidative, inflammatory, and lipoprotein alterations, with miR-23a-5p emerging as promising biomarkers and potential regulators of redox and lipid homeostasis in atherogenesis.

## 1. Introduction

Cardiovascular disease (CVD) comprises a spectrum of disorders affecting the heart and vascular system and remains the leading cause of global morbidity and mortality. According to the World Health Organization, CVD accounts for approximately 17.9 million deaths annually, underscoring the imminent necessity for more effective strategies for early diagnosis, prevention, and intervention [1,2]. Atherosclerosis, the pathological hallmark of most CVDs, is a chronic, progressive inflammatory condition which is characterized by the accumulation of lipids, immune cell infiltration, and fibrotic remodeling of the arterial wall accompanied with major clinical manifestations such as myocardial infarction, ischemic stroke, and peripheral artery disease [3,4] contributing significantly to both mortality and long-term disability [5,6]. Initiation of atherogenesis involves endothelial dysfunction and lipid oxidation, specifically apoB100 containing low-density lipoprotein (LDL) in the intima, that triggers recruitment of circulating monocytes. These monocytes differentiate into macrophages and try to limit the accumulation of oxLDL via the scavenging receptor CD36 to maintain vascular homeostasis. Through uptake of oxidized LDL (oxLDL), macrophages are transformed into foam cells which is defined as a critical event in early plaque development [7–9]. While lipid panels remain standard for assessing cardiovascular risk, their predictive accuracy for subclinical atherosclerosis and plaque vulnerability is limited. Moreover, the unpredictable nature of plaque rupture and the silent evolution of early lesions highlight the need for dynamic molecular markers that can reflect underlying cellular processes in real time [10,11]. OxLDL, the initial trigger associated with early atherosclerotic plaque formation [12,13] has been ignored as a diagnostic biomarker. On the other hand, MicroRNAs (miRNAs) have emerged as a promising molecular tool for diagnosis and prognosis. These small (∼22 nucleotide), non-coding RNAs regulate gene expression by binding to complementary sequences in the 3′ or 5′ untranslated regions (UTRs) of messenger RNAs (mRNAs), leading to translational repression or mRNA degradation [14,15]. Approximately 60% of human protein-coding genes are estimated to be regulated by miRNAs, underscoring their central role in maintaining cellular homeostasis. They have been implicated in nearly all key molecular processes driving arterial remodeling and atherosclerotic progression, including endothelial dysfunction, monocyte adhesion, vascular smooth muscle cell (VSMC) activation, and plaque formation [16]. Recent studies highlight miRNAs as both regulators and biomarkers of cardiovascular pathology. Winski et al. (2025) suggested that the clinical potential of circulating miRNAs in abdominal aortic aneurysm (AAA), a condition for which no validated biomarker currently exists [17]. Similarly, Rao et al. (2024) reviewed the role of miRNAs in diabetic macroangiopathy, revealing their involvement in vascular dysfunction associated with metabolic stress [18].

The temporal dynamics of miRNAs under high-cholesterol dietary exposure to rats and their correlation with key inflammatory, oxidative stress and lipid biomarkers have not been investigated. To fill this knowledge gap, a longitudinal high-cholesterol diet (HCD) rodent model was employed to (1) map the expression dynamics of key miRNAs during diet-induced atherogenesis, (2) correlate these miRNAs with molecular markers of vascular injury and oxidative stress (3) assess their potential as early biomarkers in cardiovascular disease.

## 2. Materials and methods

### 2.1 Source of chemicals

Cholesterol and Cholic acid were purchased from Sisco Research Laboratories Pvt. Ltd., India (SRL). Malondialdehyde (MDA) and RNAlater® were purchased from Sigma Aldrich, USA. miRNeasy serum/plasma kit purchased from Qiagen, Germany. SYBR™ Green Universal Master Mix from Applied biosystems,USA. iScript cDNA synthesis kit and Clarity Western ECL Substrate kit from BioRad, Hercules, CA, USA. miRNA, stem loop primers were purchased from Bioserve, Hyderabad, India. Primary and secondary antibodies were procured from Abcam antibodies, USA; Cell Signaling technologies, USA; Thermoscientific, USA and Abclonal, China. All other chemicals used were of analytical grade, obtained from Sigma Aldrich,USA; Medox Biotech India Pvt. Ltd., India (Medox); Sisco Research Laboratories Pvt. Ltd., India (SRL).

### 2.2 Animal studies

Age-matched (8 weeks old) Wistar male albino rats weighing around 200 ± 10 g were procured from Central Animal House Facility, University of Madras, Taramani Campus, Chennai (Tamilnadu). The animal experiments were carried out as per the guidelines of the Committee for the Purpose of Control and Supervision of Experiments on Animals (CPCSEA), India and approved by the Institutional Animal Ethics Committee (IAEC), University of Madras, Tamilnadu (IAEC Approval No. 01/03/2023).

**Figure 1.**
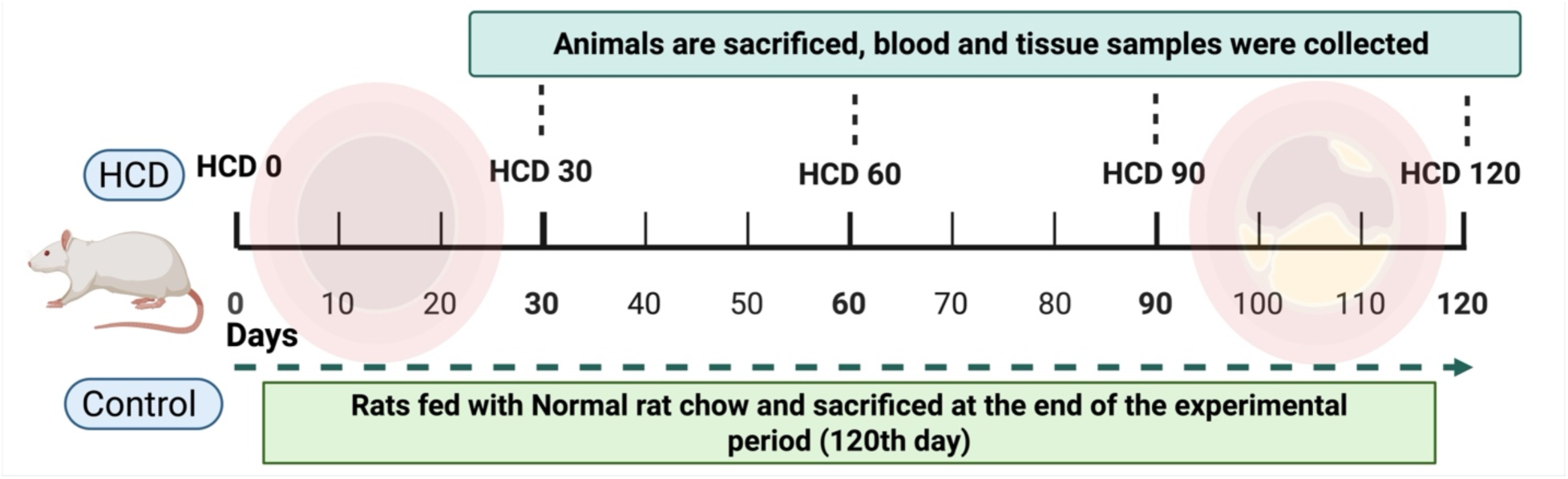
Experimental groups

Rats were acclimatized for one week under standard laboratory conditions and randomization were carried out before initiating the experiment. High cholesterol diet (HCD) model was established by feeding High-cholesterol Diet (comprising of the normal rat chow supplemented with 4% cholesterol and 1.5% cholic acid) till the end of the experimental period. At the end of the experimental period (30, 60, 90 and 120days), rats were anesthetized with ketamine (22mg/kg body weight, i/p). Aorta was excised, washed with ice-cold physiological saline, and stored at -80 °C for biochemical and molecular studies. For histopathological studies, aortic tissue was immersed in 4% paraformaldehyde for 48 hours and transferred to PBS for further analysis.

### 2.3 Lipid peroxidation

Malondialdehyde levels in serum were measured as described by Devasagayam and Tarachand (1974)

### 2.4 miRNA isolation

miRNA was isolated using miRNeasy serum/plasma kit (Qiagen, Germany) according to the manufacturer’s instructions. Then the eluted miRNA was quantified using NanoDrop2000 UV– Vis spectrophotometer (Thermo Scientific, USA)

#### 2.4.1 cDNA synthesis and Real-Time PCR

The cDNA conversion was carried out from miRNA using custom designed miRNA specific stem-loop primers for miRNAs (Table 1) and 10 ng of RNA for studying miRNA expression. The miRNA samples were pre-incubated at 65 °C for 20 min followed by 55 °C for 90 min, 72 °C for 15 min and final hold at 4 °C. cDNA conversion was performed using a Reverse Transcription kit (Biorad, USA) and cDNAs were further diluted 25-fold and stored at − 20 °C. Real-time-qPCR was performed in ABI Quantstudio 3 (ABI Lifetechnology, USA). The reactions were performed in 96 well optical plates with 10 μL total volume using 1 μL of cDNAs as template, 5μL of SYBR green 2X Universal Master mix (Biorad, Herculus, USA), 0.4μL of specific forward primers and 0.4 μL of universal reverse primer (Table 1) by following the thermal cycling conditions: 50 °C for 2 min and 95 °C for 10 min once, followed by 95 °C for 15 s and 60 °C for 1 min for 40 cycles to determine miRNAs expression. miR-423-5p was used as an endogenous control. NTC reactions were included in all the experiments. All the reactions were carried out in triplicates; mean Ct was used for analysis and the expression level was calculated using 2^−ΔΔCt^ method.

**Table 1:**
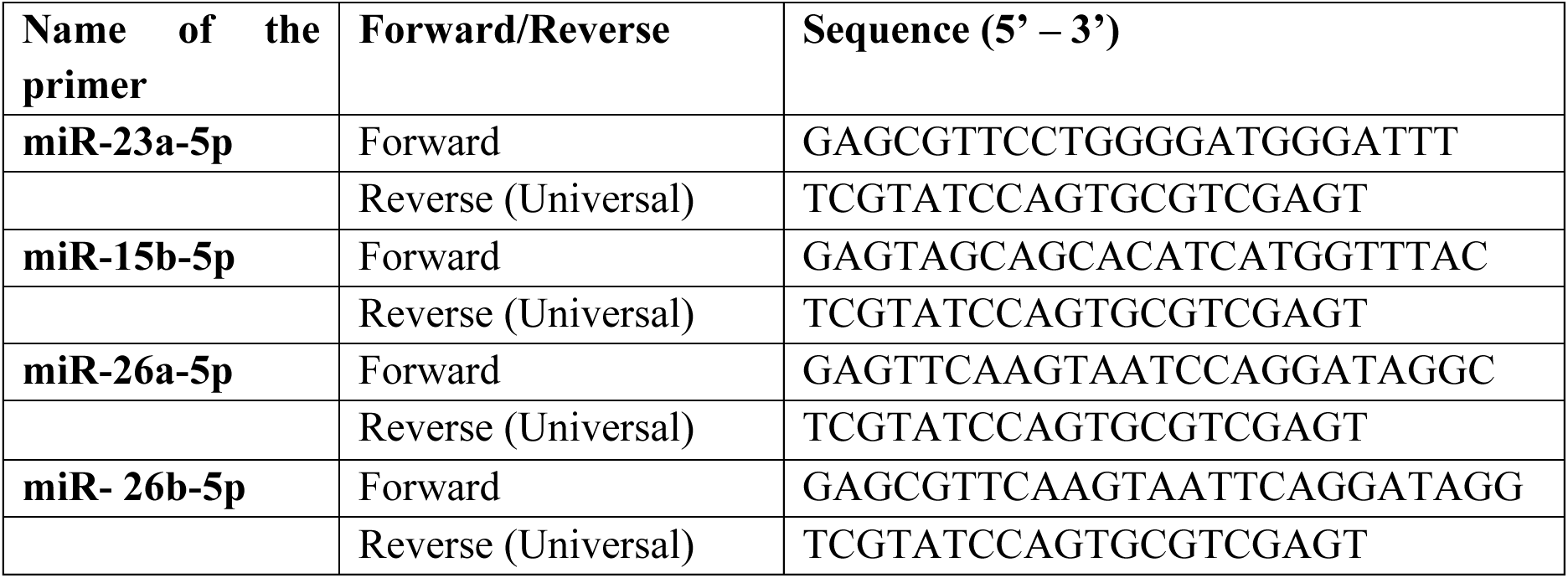
miRNA forward and reverse sequences were used for performing RT PCR.

### 2.5 Bioinformatic studies on miRNA targets

Target scan, miRDB, miRTarbase are the databases used to verify the microRNAs and its targets. We also explored the pathways using the KEGG database that could be modulated by the miRNA expression and were then the expressions were measured.

### 2.6 ELISA

Serum levels of Apolipoprotein A1, Paraoxonase 1, oxLDL, inflammatory cytokines (CRP, TNF-α, IL-6 and IL-8) and cell adhesion molecules (VCAM1,) were assessed in all experimental groups using ELISA. Briefly, 25 μg of serum protein was coated onto 96-well plates using bicarbonate buffer and incubated overnight at 4°C. The following day, the plates were washed with 1X PBS and blocked with 1% BSA solution (200 μl/well) for 2 hours to prevent non-specific binding. After washing the wells thrice with 1X PBS and allowing them to dry, respective primary antibodies diluted in 1% BSA (1:1000), were added (100 μl/well) and incubated overnight at 4°C. The next day, the wells were washed four times with PBS-Tween-20 and incubated with horseradish peroxidase-conjugated secondary antibodies for 1 hour at room temperature. After additional washes, 100 μl of ABTS (2,2’-Azino-bis (3-ethylbenzothiazoline-6-sulfonic acid)) substrate was added and the plates were incubated in the dark for 20 minutes. To stop the reaction, 100 μl of 2 N sulfuric acid was added to each well, and the absorbance was measured at 415 nm using a Bio-Rad iMark™ Microplate Absorbance Reader (Bio-Rad Laboratories, Hercules, CA, USA).

### 2.7 Immunoblotting

50ug of protein from rat aortic tissues were subjected to 6-12% polyacrylamide gel electrophoresis and then transferred onto a polyvinylidene difluoride membrane (Biorad, USA). Membranes were blocked with 5% skimmed milk in Tris-buffered saline with 0.1% Tween 20 for 1 h and then incubated overnight with primary antibody diluted with 0.8% bovine serum albumin at 4°C. Antibodies specific for APOA1, PON1, ABCA1, ABCG1, CD36, Alpha SMA, oxLDL and NFkB were used. β-Actin served as a loading control. The samples were then incubated with the appropriate secondary antibodies conjugated to horseradish peroxidase. Signals were detected using enhanced luminescence and the intensity of protein bands was quantified using ImageJ software.

### 2.8 Human Studies

The present study was conducted after the approval by the Institutional Ethics Committees (IEC) of Sri Ramachandra Medical College and Hospitals and within the ethical framework of Dr. ALM PG Institute of Basic Medical Sciences, University of Madras, Chennai. The blood samples were collected from apparently healthy individuals and patients with CAD. The study includes a total of 20 participants diagnosed with CAD and 20 healthy controls. The patient’s contextual and clinicopathological characteristics were documented with standard questionnaire following the IEC guidelines and written informed consent was obtained from each patient, after explaining about the research study. The blood samples were collected and centrifuged 3000rpm for 15minutes for serum separation and transported to the laboratory in cold storage and stored at – 80 °C for miRNA isolation.

#### 2.8.1 miRNA isolation cDNA synthesis and RT PCR

The procedure carried out was previously described in the above sections (Section 3.4)

### 2.9 Statistical analysis

Data were analyzed using Graph Pad Prism statistical software, v 8.01 (Graph Pad software Inc, USA). Univariate or factorial analysis of variance (ANOVA) with the Tukey post hoc test was used. Values are expressed as Mean ± SD, where * represents (p < 0 05), ** represents (p < 0.01) and *** (p < 0 001).

## 3. Results

### 3.1 High-cholesterol diet induces dyslipidemia and elevates atherogenic risk in a time-dependent manner

To assess the metabolic impact of prolonged high-cholesterol diet (HCD) exposure, serum lipid profiles and atherogenic index (AI) were evaluated at 30, 60, 90, and 120 days. A progressive dyslipidemic shift was observed on HCD feeding. Total cholesterol (TC) levels increased significantly over time, rising from 123.45 ± 9.1 mg/dL in controls to 131.45 ± 8.0 mg/dL at 30 days, and further to 157.60 ± 13.4 mg/dL (p < 0.01), 175.38 ± 13.7 mg/dL (*p* < 0.001), and 202.15 ± 17.6 mg/dL (*p* < 0.001) at 60, 90, and 120 days, respectively. Triglyceride (TGL) levels followed a similar trajectory, increasing from 100.76 ± 6.21 mg/dL in controls to 174.32 ± 5.10 mg/dL at 120 days (*p* < 0.001), with moderate increase at 30, 60, and 90 days.

VLDL concentrations rose significantly from 20.14 ± 1.71 mg/dL in controls to 34.29 ± 2.31 mg/dL at 120 days (*p* < 0.001), while LDL levels nearly doubled, reaching 149.65 ± 6.95 mg/dL at 120 days compared to 70.01 ± 4.20 mg/dL in controls (*p* < 0.001). In contrast, HDL levels declined progressively, from 34.94 ± 2.61 mg/dL in controls to 19.89 ± 2.5 mg/dL at 120 days (*p* < 0.001), indicating a shift towards a more atherogenic lipid profile.

Consistent with these alterations, the atherogenic index (AI) increased markedly with HCD duration, rising from 0.1002 in controls to 0.486 at 120 days. These findings collectively demonstrate that chronic HCD exposure induces a robust dyslipidemic response characterized by elevated pro-atherogenic lipoproteins, reduced HDL, and a significantly increased cardiovascular risk profile.

### 3.2 Progressive elevation of serum lipid peroxidation reflects time-dependent oxidative stress induced by high-cholesterol diet

Serum lipid peroxidation (LPO), a key indicator of oxidative stress, was significantly elevated in rats subjected to high-cholesterol diet (HCD) feeding. Compared to the control group, which exhibited baseline LPO levels of approximately 50 µmol/mg protein, HCD-fed rats demonstrated a time-dependent rise in LPO. After 30 days of HCD, LPO levels nearly doubled (∼100 µmol/mg protein, *p* < 0.05), and continued to increase at 60 days (∼150 µmol/mg protein, p < 0.01) and 90 days (∼175 µmol/mg protein, *p* < 0.001). The most pronounced oxidative response was observed at 120 days, with LPO levels reaching ∼325 µmol/mg protein (*p* < 0.001 vs. all groups). These findings indicate that prolonged HCD exposure induces a robust and progressive increase in systemic lipid peroxidation, underscoring the diet’s pro-oxidant potential.

### 3.3 High-cholesterol diet disrupts antioxidant defense and promotes lipoprotein oxidation

To investigate the oxidative and lipoprotein-modifying effects of prolonged high-cholesterol diet (HCD) exposure, serum levels of paraoxonase-1 (PON1), apolipoprotein A1 (ApoA1), and oxidized LDL (oxLDL) were quantified across experimental groups. PON1, a key antioxidant enzyme associated with HDL, was significantly reduced in all HCD-fed groups compared to controls. Control rats exhibited the highest PON1 levels (∼130 pg/mL), while HCD30, HCD60, HCD90, and HCD120 groups showed a marked and progressive decline, with the most pronounced reduction observed at 120 days (*p* < 0.001).

Similarly, ApoA1 levels, a major structural and functional component of HDL, were significantly diminished in HCD-fed rats. Control animals displayed ApoA1 concentrations of approximately 200 pg/mL, which declined significantly across all HCD groups, with statistical significance ranging from *p* < 0.05 to p < 0.01, indicating impaired HDL functionality.

In contrast, oxLDL levels, a marker of lipid peroxidation and atherogenic risk were significantly elevated in HCD-fed rats. While control animals maintained low oxLDL levels (∼50 pg/mL), HCD60, HCD90, and HCD120 groups exhibited a robust increase (*p* < 0.001), consistent with enhanced oxidative modification of LDL. These findings collectively suggest that chronic HCD feeding compromises antioxidant defenses and promotes lipoprotein oxidation, contributing to a pro-atherogenic and pro-oxidative systemic environment.

### 3.4 High-Cholesterol Diet Induces Systemic Inflammation and Endothelial Activation

To evaluate the pro-inflammatory and vascular effects of prolonged high-cholesterol diet (HCD) exposure, circulating levels of C-reactive protein (CRP), tumor necrosis factor-alpha (TNF-α), interleukin-6 (IL-6), vascular cell adhesion molecule-1 (VCAM-1), and interleukin-8 (IL-8) were quantified across experimental groups. A time-dependent increase in all measured parameters was observed, indicating progressive systemic inflammation and endothelial dysfunction.

CRP levels, a sensitive marker of systemic inflammation, increased from ∼50 ng/mL in control rats to ∼160 ng/mL in the HCD120 group (*p* < 0.001), with intermediate elevations at 30, 60, and 90 days. TNF-α levels followed a similar trend, rising from ∼40 pg/mL in controls to ∼70 pg/mL at 120 days (*p* < 0.001), reflecting enhanced pro-inflammatory cytokine activity.

IL-6 levels exhibited a marked increase, from ∼20 pg/mL in controls to ∼65 pg/mL in the HCD120 group (*p* < 0.001), with significant elevations observed as early as 60 days. VCAM-1, a key adhesion molecule involved in leukocyte recruitment and endothelial activation, inclined sharply from ∼40 ng/mL in controls to ∼170 ng/mL at 120 days (*p* < 0.001), indicating progressive vascular inflammation.

IL-8 levels also increased steadily, from ∼70 pg/μL in controls to ∼95 pg/μL at 90 days, suggesting enhanced chemotactic signaling and neutrophil activation. Collectively, these findings demonstrate that chronic HCD feeding elicits a robust inflammatory response and promotes endothelial activation, contributing to a pro-atherogenic and pro-oxidative systemic environment.

### 3.5 Impact of High-Cholesterol Diet on Protein Expression in Animal Models

Densitometric analysis of Western blot bands was performed to quantify the expression levels of Alpha Smooth Muscle Actin (Alpha SMA), ATP-binding cassette sub-family G member 1 (ABCG1), and ATP-binding cassette sub-family A member 1 (ABCA1) in rat aortic tissues across four experimental groups. Alpha SMA expression was highest in the control group and showed a progressive and statistically significant decrease with increasing treatment duration. HCD60 exhibited a reduced expression compared to control (p < 0.05), while HCD90 demonstrated a more pronounced reduction (p < 0.01). The lowest expression was observed in HCD120 (p < 0.001), indicating a strong time-dependent downregulation. In contrast, ABCG1 levels were significantly lower in HCD60 compared to control (p < 0.05), but no further significant changes were observed in HCD90 or HCD120 (both p > 0.05) relative to HCD60. This suggests an early suppression of ABCG1 expression which plateaued with extended treatment durations. Similarly, ABCA1 expression declined significantly in the HCD60 group versus control (p < 0.05) and remained statistically unchanged in HCD90 (p > 0.05). However, a further significant reduction was observed in HCD120 compared to HCD90 (p < 0.01), pointing toward progressive suppression of cholesterol transport machinery with prolonged treatment.

The expression of CD36, NFκB, and oxLDL in rat aortic tissue under varying durations of high-cholesterol diet (HCD). CD36, a scavenger receptor linked to lipid uptake and foam cell formation, showed a progressive and statistically significant increase across HCD groups. Compared to control, HCD60 rats exhibited elevated levels of CD36 (p < 0.05), with further enhancement in HCD90 (p < 0.01) and a pronounced peak in HCD120 (p < 0.001), suggesting cumulative lipid-driven macrophage activation. NFκB, a central transcription factor mediating inflammatory signaling, mirrored CD36 expression trends. NFκB levels were significantly upregulated beginning at HCD60 (p < 0.05), intensifying at HCD90 (p < 0.01) and reaching maximal levels in HCD120 (p < 0.001), indicating progressive endothelial inflammation and pro-atherogenic activation. oxLDL, an atherogenic lipid marker, was likewise elevated in response to HCD exposure. A modest increase was noted in HCD60 (p < 0.05), which intensified in HCD90 (p < 0.01) and peaked in HCD120 (p < 0.001). This accumulation reflects enhanced lipid peroxidation and vascular oxidative stress. Together, the rising levels of CD36, NFκB, and oxLDL in HCD groups underscore the presence of strong chronic inflammation and oxidative injury associated with diet-induced atherosclerosis progression in rat aortic tissue. These findings reinforce the link between lipid overload, inflammatory activation, and vascular pathology.

### 3.6 miR-23a-5p and miR-15b-5p Are Differentially Expressed in Response to High-Cholesterol Diet

To explore the potential of circulating microRNAs as biomarkers of high-cholesterol diet (HCD)-induced pathophysiology, the expression levels of miR-23a-5p and miR-15b-5p were quantified using qRT-PCR and analyzed via the 2^−ΔΔCT method across experimental groups. miR-23a-5p expression remained relatively stable during the early phases of HCD exposure, with modest increases at 30 and 60 days (∼1.5- to 2-fold vs. control), but without statistical significance. However, a marked and statistically significant upregulation was observed at 120 days, with expression levels reaching ∼4.8-fold compared to control (*p* < 0.001), suggesting a late-stage response to prolonged dietary stress. In contrast, miR-15b-5p exhibited an earlier and more dynamic expression pattern. A significant increase was detected as early as 60 days (∼4.3-fold, *p* < 0.05), followed by sustained elevation at 90 and 120 days (∼3- to 4-fold, *p* < 0.05 for HCD120). These findings indicate that miR-15b-5p may serve as an earlier marker of HCD-induced molecular alterations, while miR-23a-5p may reflect more advanced or cumulative effects of dietary cholesterol. Together, these data highlight the differential temporal regulation of miR-23a-5p and miR-15b-5p in response to HCD and support their potential utility as circulating biomarkers for monitoring disease progression and oxidative stress-related vascular dysfunction.

### 3.7 miR-23a-5p Expression Correlates with Oxidative and Lipoprotein Biomarkers

To explore the potential mechanistic relevance of miR-23a-5p in high-cholesterol diet (HCD)-induced oxidative stress and lipoprotein dysfunction, correlation analyses were performed between miR-23a-5p expression (2^−ΔΔCt) and key circulating biomarkers, oxidized LDL (oxLDL), apolipoprotein A1 (ApoA1), and paraoxonase-1 (PON1). A strong positive correlation was observed between miR-23a-5p and oxLDL levels (R²=0.5393), suggesting that elevated miR-23a-5p expression is associated with increased lipid peroxidation and oxidative modification of LDL. In contrast, miR-23a-5p expression was inversely correlated with ApoA1 levels (R²=0.2738) and PON1 levels (R²=0.3294), indicating a potential link between miR-23a-5p upregulation and impaired HDL functionality and antioxidant defense. These findings support the hypothesis that miR-23a-5p may serve not only as a biomarker of oxidative stress and lipoprotein imbalance but also as a potential regulator of redox-sensitive pathways in the context of HCD-induced vascular dysfunction.

### 3.8 miR-23a-5p Expression in healthy control and CAD subjects

Assessment of miR-23a-5p expression revealed a significant upregulation in CAD subjects compared to control individuals, as shown in densitometric measurements. The relative expression level of miR-23a-5p was markedly higher in the CAD group (p < 0.01), indicating a potential association with the pathophysiology of coronary atherosclerosis. Elevated miR-23a-5p levels in CAD patients reflect its possible role in promoting vascular inflammation, endothelial dysfunction, and smooth muscle proliferation which is hallmark features of atherosclerotic plaque development. Prior evidence suggests that miR-23a-5p may influence NFκB-mediated transcription, contribute to monocyte adhesion, and impair cholesterol efflux mechanisms, thereby accelerating the inflammatory cascade seen in atherosclerotic lesions. These findings support the hypothesis that miR-23a-5p is a pro-atherogenic microRNA and may serve as both a biomarker for early atherosclerotic changes and a therapeutic target in coronary artery disease.

## 4.0 Discussion

The asymptomatic process of atherosclerosis is initiated by lipid retention in the intima, fatty streak formation, and plaque development. This concealed process comes into limelight when symptoms emanate in an eventuality such as a heart attack or stroke. Lipoprotein measurements deem an individual to be atherosclerotic prone but neither dictate under which stage the events are set in the arteries nor the efficacy of the therapeutic strategies. Research into the crucial role of miRNAs in the formation of plaques and rupture is still in its infancy and therefore work is required to pinpoint the exact miRNA panel that can be employed as a biomarker. There is a need for newer therapeutic targets and combinatorial approaches for the management of subclinical and clinical atherosclerosis because interventions to lower LDL or boost HDL do not ensure cessation of plaque progression or rupture.

The pronounced elevations in total cholesterol, triglycerides, LDL-C, and VLDL-C, accompanied by an evident decrement of HDL-C and a heightened atherogenic index, mirror the pathological lipid perturbations observed in the current study which is a characteristic feature of diet-induced atherogenesis (19, 20). Dyslipidemia, oxidative stress and inflammation considered as dreadly traids of atherosclerosis. Given the central role of oxidative stress in this process, assessing systemic redox balance through biomarkers such as serum lipid peroxides and antioxidant capacity is essential. In the present study, a significant increase in serum lipid peroxides was observed in HCD fed animals. These findings are consistent with the previous reports (21) highlighting the role of lipid peroxidation in the pathogenesis of atherogenesis. The observations made in HCD fed rats assure that dyslipidemic aberrations have been further compounded by a substantial increase in lipid peroxidation, underscoring the pro-oxidative milieu instigated by a hypercholesterolemic diet. Elevated malondialdehyde (MDA)concentrations corroborate the existence of severe oxidative derangement, as recorded in both preclinical models and dyslipidemic cohorts (22, 23). Further elevated ROS levels contribute to endothelial dysfunction and promote the oxidative modification of low-density lipoproteins (ox-LDL), a key step in foam cell formation and plaque development (24, 25). Brown and Goldstein were the first to postulate that LDL undergoes some structural modifications to achieve atherogenic properties (26, 27). OxLDL is inclined in the groups of animals fed with HCD when compared to control which reflects the foam cell formation in the arteries.

Paraoxonase, an enzyme associated with HDL is involved in protecting LDL from oxidation (28) and so the level of PON is measured in the various experimental groups. The PON 1 levels and activity were reduced in the HCD fed animal when compared to the control animals. Similar kind of observations denoting a diminution in PON level has been shown to be associated with increased oxidative stress in serum and coronary events (29). This ascertains the inverse relationship between PON and ox-LDL and emphasizes the importance of PON as an anti-atherogenic protein. Apolipoprotein A1 (ApoA1), the major structural protein of HDL, plays a central role in reverse cholesterol transport and antioxidant defense. The mechanistic underpinning of the cardioprotective effect of ApoA1 is further bolstered by a seminal report in Cochran et al., (2021), which revealed an inverse association between cholesterol efflux capacity and cardiovascular events, independent of HDL cholesterol levels highlighting the function of ApoA1. Moreover, research from the JUPITER trial reported that HDL particle number and cholesterol efflux capacity is strongly linked to ApoA1 concentration and it is a predictive marker of incident cardiovascular events, even independent of HDL-C levels (30, 31). In animal models, ApoA1 deficiency has been associated with increased plaque burden, yet HDL/ApoA1 replacement therapy reduces lesion area, reinforcing its atheroprotective role (32). Prolonged high-cholesterol diet (HCD) exposure, such as over 120 days, has been shown to significantly reduce ApoA1 levels in experimental animal models when compared to control groups. This decline is critical, as it correlates with impaired cholesterol efflux and increased susceptibility to lipid accumulation and vascular inflammation.

Further, Reactive oxygen species (ROS) play a pivotal role in driving the chronic inflammatory cascade (33). Status of inflammation was assessed in all the experimental groups with reference to TNF-α, CRP, IL6 and IL8 as levels of CRP that are consistently associated with cardiovascular diseases particularly myocardial infarction (34). Pro-inflammatory and inflammatory cytokines such as Tumor necrosis factor-alpha (TNF-α), interleukin-6 (IL-6), C-reactive protein (CRP), and interleukin-8 (IL-8) exhibit a progressive increase in animals fed with a high-cholesterol diet (HCD) with levels significantly elevated by day 60 and peaking at days 90 and 120 compared to control groups. This temporal incline in pro-inflammatory cytokines underscores the advancing inflammatory milieu and supports the progression of atherosclerotic pathology and recent studies have also showed the same trend in patients (35). TNF-α activates the gene expression of diverse inflammatory cytokines and chemokines, either directly or indirectly through the activation of various transcriptional factors like NF-κB and promotes cell death (36). Several studies also validate the involvement of CRP as a potent mediator of atherogenesis contributing to the advancement of the atherosclerotic lesion through a direct pro-inflammatory effect. CRP brings about the release of endothelin-1; up regulates the expression of adhesion molecules and other chemokines in the endothelial cell layers and vascular smooth muscle cells are implicated to occupy a prominent place in the process of atherosclerosis with the inflammatory cells playing a prime role in the recruitment of adhesion molecules. VCAM-1 levels show a marked elevation in rats fed a high-cholesterol diet (HCD) for 90 days, with peak expression observed at 120 days, indicating progressive endothelial activation associated with atherosclerotic development.

Smooth muscle cells (SMCs) play dual roles in atherosclerosis by contributing to lesion initiation, foam cell formation, and inflammation, while also stabilizing plaques through fibrous cap formation. Their phenotypic plasticity allows them to shift between pro-atherogenic and protective states. This dynamic behavior makes SMCs key regulators of plaque progression and stability (37). In rats fed a high-cholesterol diet (HCD), α-SMA expression was progressively reduced at both 90 and 120 days compared to control animals, suggesting a phenotypic shift of vascular smooth muscle cells from a contractile to a synthetic state which is an early hallmark of vascular remodeling and atherogenesis. miR-15b-5p expression displayed a biphasic pattern in rats subjected to a high-cholesterol diet (HCD), with elevated levels at days 60 and 120, and a transient dip at day 90 though still above baseline. The transient reduction at HCD90 could indicate a compensatory phase or cellular adaptation, while the re-elevation at HCD120 may correspond to advanced lesion development and vascular remodeling. Recent studies support the multifaceted role of miR-15b-5p in cardiovascular pathology. For instance, Martino et al. (2023) (38) demonstrated that miR-15b-5p promotes endothelial dysfunction by targeting SIRT4, contributing to oxidative stress and inflammation in vascular tissues. Moreover, miR-15b-5p has been shown to influence angiogenesis and arteriogenesis by targeting AKT3, further linking it to atherosclerotic progression (39, 40). Hence the circulatory miR-15b-5p might not indirectly reflect the inhibition of collateral formation. However, evidence from the human studies state that despite an adaptive mechanism in miR-15b-5p tends to downregulate in CAD, poor collaterals are formed. The present study demonstrates that prolonged exposure to a high-cholesterol diet (HCD) for 120 days leads to a significant downregulation of ATP-binding cassette transporters A1 and G1 (ABCA1 and ABCG1) in rats, accompanied by an upregulation of miR-23a-5p. These findings suggest a mechanistic link between microRNA-mediated post-transcriptional regulation and impaired cholesterol efflux capacity in the context of diet-induced dyslipidemia. ABCA1 and ABCG1 are pivotal in facilitating reverse cholesterol transport by promoting cholesterol efflux to apolipoprotein A-I and HDL, respectively. Their diminished expression under HCD conditions likely contributes to intracellular lipid accumulation and foam cell formation, accelerating atherogenesis.

The observed increase in miR-23a-5p expression may play a regulatory role in this process. Bioinformatic predictions and experimental studies have identified ABCA1 and ABCG1 as potential targets of miR-23a-5p, implicating this microRNA in the suppression of cholesterol transporter expression. This aligns with previous reports indicating that miR-23a-5p is upregulated in hyperlipidemic states and may contribute to lipid dysregulation and vascular inflammation. The concurrent elevation of miR-23a-5p and suppression of ABCA1/ABCG1 in HCD-fed rats supports the hypothesis that miRNA-mediated silencing mechanisms are involved in the progression of atherosclerotic changes.

A progressive upregulation of miR-23a-5p and concurrent downregulation of ABCA1, ABCG1 observed in HCD-fed animal models, on feeding 120days, an increase in miR-23a-5p is observed. Even though further experiments are warranted to identify whether it is a cause or effect, the study emphasizes that circulatory miR-23a-5p is shown to correlate with the atherosclerotic plaque progression. we extended our investigation to human serum samples. Specifically, we assessed the expression of circulating miR-23a-5p in individuals with coronary artery disease (CAD) compared to healthy controls to evaluate its potential as a non-invasive biomarker of atherosclerotic burden.

In this study, we observed a significant upregulation of circulating miR-23a-5p in the serum of patients with coronary artery disease (CAD) compared to healthy controls. This finding aligns with emerging evidence suggesting that miR-23a-5p plays a pivotal role in cardiovascular pathology through its regulation of lipid metabolism, inflammation, and endothelial function. Elevated levels of miR-23a-5p have been previously reported in patients with heart failure and reduced ejection fraction, indicating its broader involvement in cardiovascular remodeling and systemic stress responses. Mechanistically, miR-23a-5p has been shown to target genes such as ABCA1 and ABCG1, which are essential for cholesterol efflux and reverse cholesterol transport. Its overexpression may therefore contribute to lipid accumulation and foam cell formation are the hallmarks of atherosclerotic plaque development. Furthermore, a recent systematic review highlighted the diagnostic potential of miRNAs, including miR-23a, in CAD and hypertension, reinforcing its utility as a non-invasive biomarker for early detection and disease monitoring (41). The consistent elevation of miR-23a-5p in both experimental and clinical settings underscores its translational relevance and warrants further investigation into its prognostic value and therapeutic modulation.

## Conclusion

This study establishes that chronic high-cholesterol diet (HCD) exposure drives a multifactorial progression of vascular dysfunction characterized by lipid metabolic imbalance, oxidative stress, and inflammatory activation. The sustained upregulation of miR-23a-5p in HCD-fed animal models, coupled with its elevated expression in CAD patients, underscores its pathological relevance in human atherogenesis. Notably, miR-23a-5p was found to interfere with lipid homeostasis through modulation of key cholesterol efflux pathways and was strongly associated with oxLDL accumulation, either as a mediator or a downstream target of lipid peroxidation. The combined metabolic and oxidative converge to impair vascular integrity, promote endothelial activation, and accelerate the development of atherosclerotic lesions. Mechanistically, miR-23a-5p was shown to influence the transcript levels of ABCA1 and ABCG1, key mediators of cholesterol efflux, suggesting its role in impairing lipid transport and homeostasis. Furthermore, a strong association was observed between miR-23a-5p and elevated oxLDL levels, indicating that miR-23a-5p either contributes to lipid oxidative modification or is induced by oxLDL-triggered signaling. These findings collectively propose miR-23a-5p as a critical molecular marker in the pathogenesis of atherosclerosis, mediating vascular dysfunction.

***Abinayaa Rajkumar***: Conceptualization (First); data curation (First); formal analysis (First); investigation (First); methodology (First); visualization (First); writing − original draft (First); writing − review and editing (First). ***Gokulprasanth Panchalingam***: Visualization and writing − review and editing (supporting). ***Deepthy Jayakumar:*** Investigation and visualization; writing − review and editing (supporting). ***Devasimman Perumal:*** Investigation and visualization (supporting)***. Pazhanisankar Muthusamy***: Investigation and visualization (supporting). ***Dhamodharan Shankar:*** Investigation and visualization (supporting). ***Divya Bhavani Ravi***: Investigation and visualization (supporting). ***Kalaiselvi Periandavan***: Conceptualization (lead); funding acquisition (lead); project administration (lead); resources (lead); supervision (lead); validation (lead); visualization (lead); writing − review and editing (lead). All authors have read and approved the final version of the manuscript

## Acknowledgements

The authors express their gratitude to the financial support provided to Abinayaa Rajkumar by **Department of Science and Technology – Science and Engineering Research Board (DST-SERB),** New Delhi, Government of India in the form of Junior Research Fellow. The authors also acknowledge **Department of Health Research - Multidisciplinary Research Unit** (DHR-MRU) at the Dr. ALM Post Graduate Institute for Basic Medical Sciences, University of Madras, for providing access to its state-of-the-art facilities.

## Conflict of Interest statement

The authors declare that they have no potential conflicts of interest, including any financial or personal relationships with other people or organizations.

## Data Availability Statement

The data supporting the findings of this study are available from the corresponding author upon reasonable request.

## Declaration of Transparency and Scientific Rigour

This Declaration acknowledges that this paper adheres to the principles for transparent reporting and scientific rigor of preclinical research recommended by funding agencies, publishers, and other organizations engaged in supporting research.

---

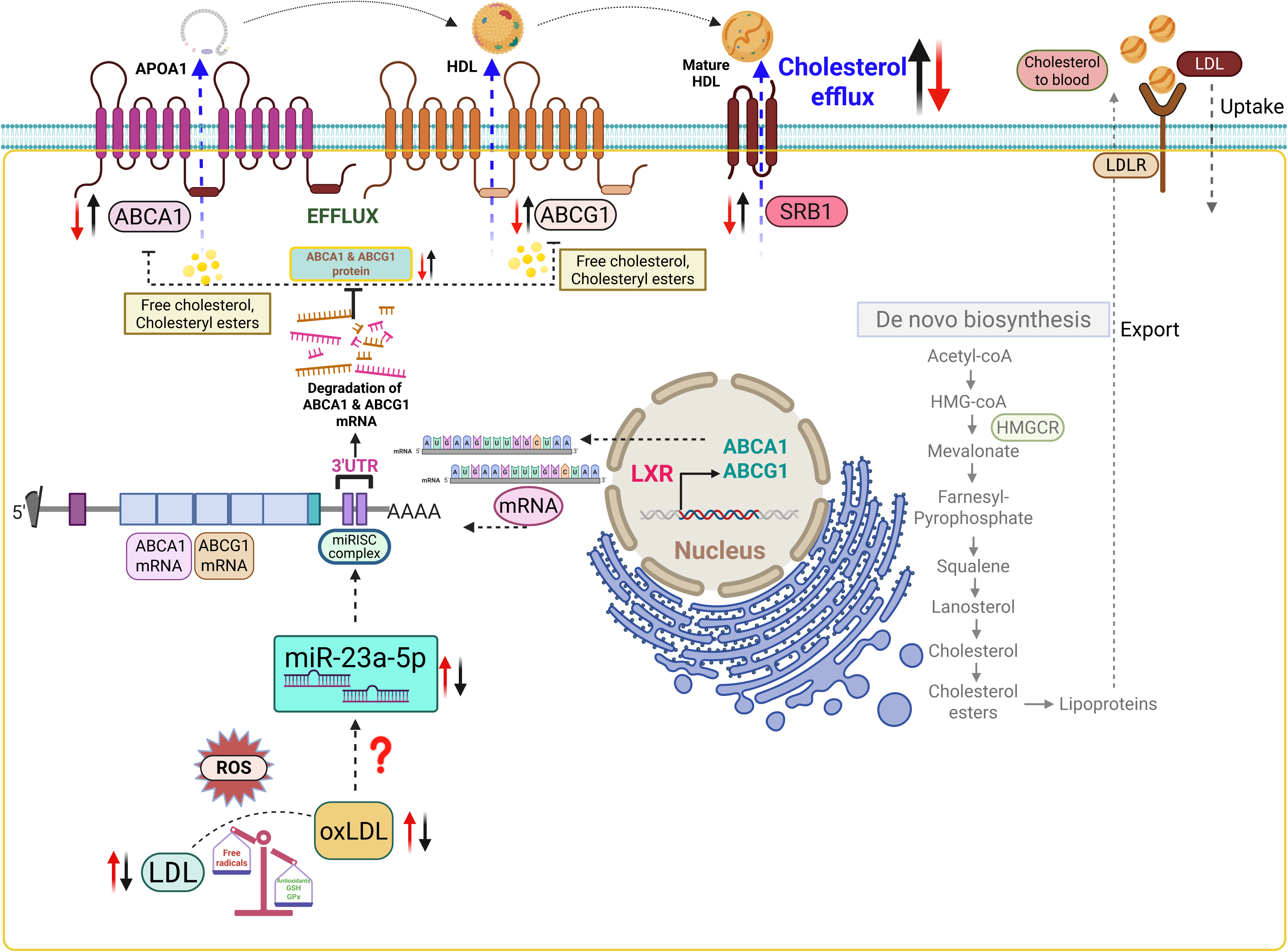

**Figure 1.**
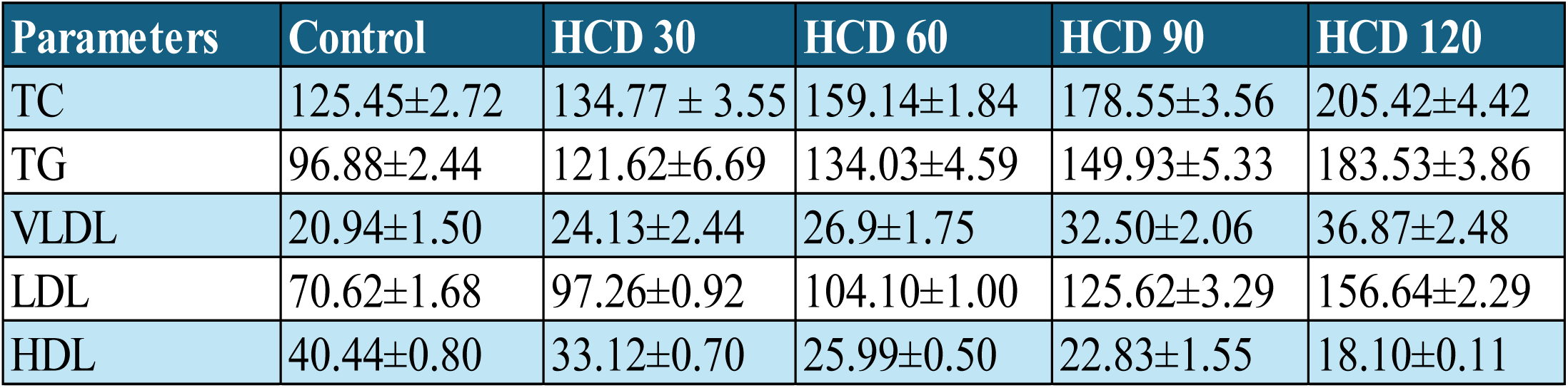
Lipid profile Values are represented as mean ± standard error of mean (SEM). Statistical analysis was performed using one-way ANOVA with Tukey’s multiple comparisons test. **\*** *p* < 0.05, **\*\*** *p* < 0.01, **\*\*\*** *p* < 0.001 vs. indicated groups.

**Figure 2.**
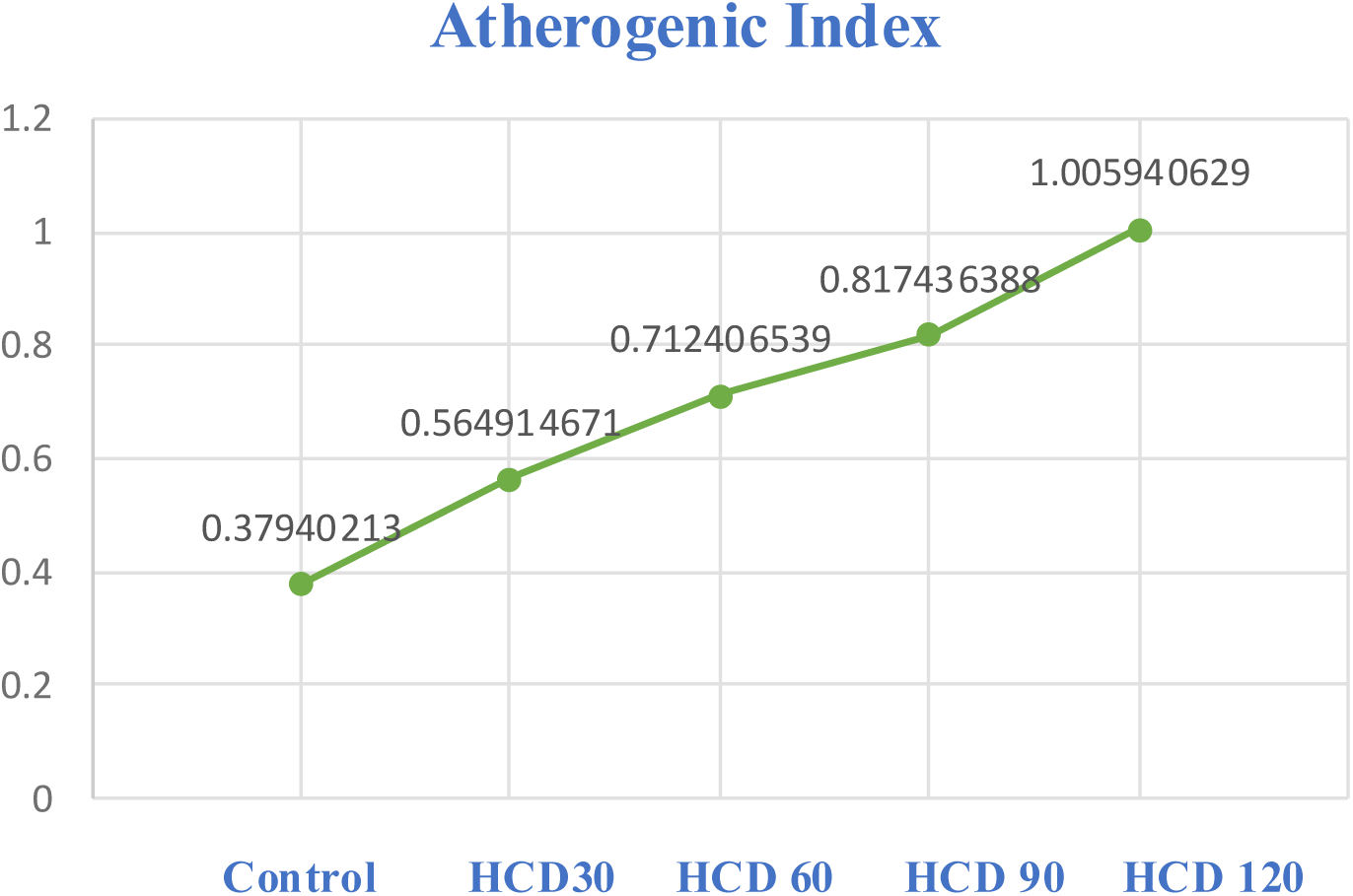
Graph depicts the Atherogenic index.

**Figure 3.**
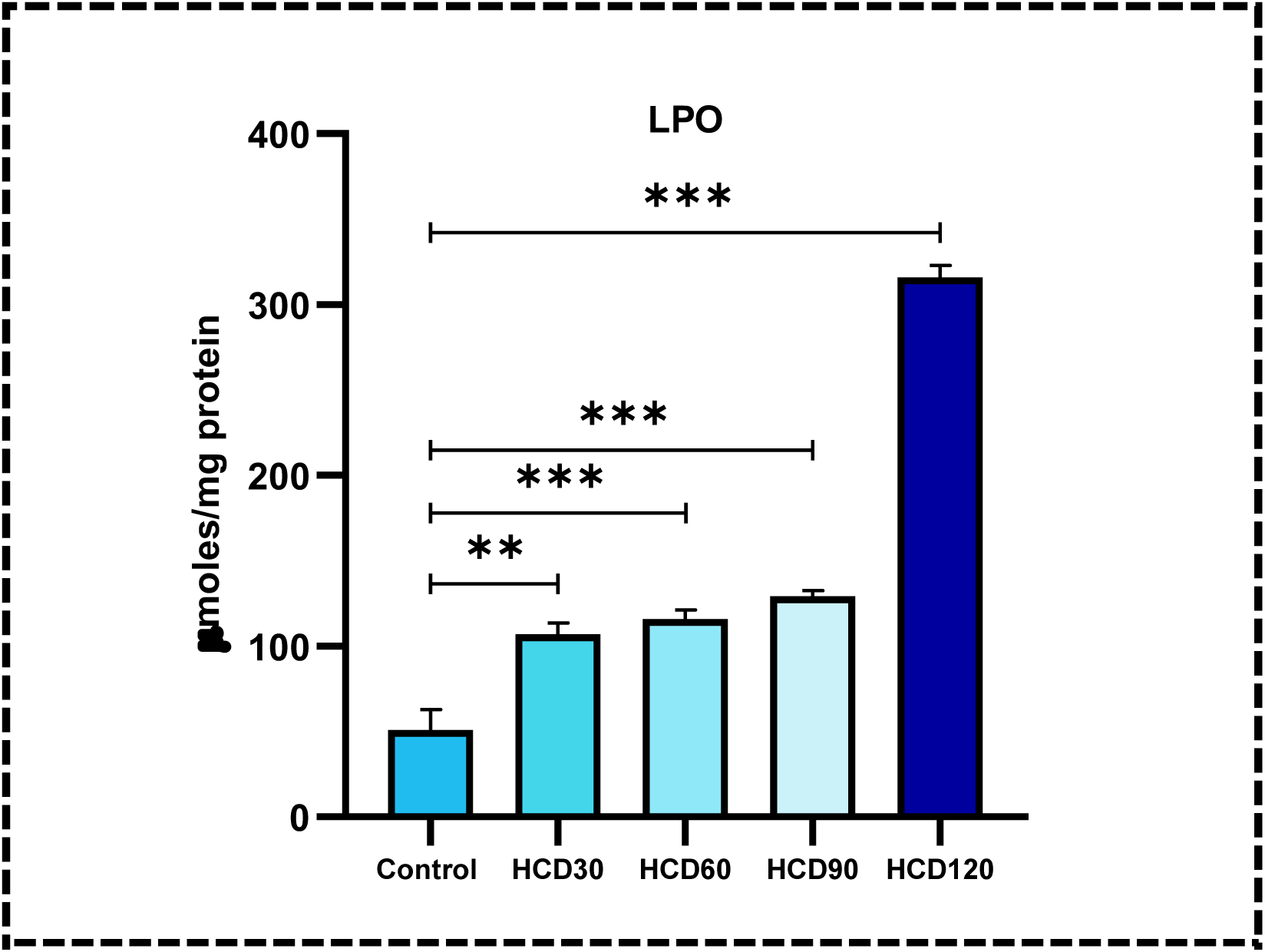
Serum lipid peroxidation (LPO) levels in rats from all experimental groups (Control and High cholesterol diet (HCD) fed rats). Values are represented as mean ± standard error of mean (SEM). Statistical analysis was performed using one-way ANOVA with Tukey’s multiple comparisons test. **\*** *p* < 0.05, **\*\*** *p* < 0.01, **\*\*\*** *p* < 0.001 vs. indicated groups.

**Figure 4.**
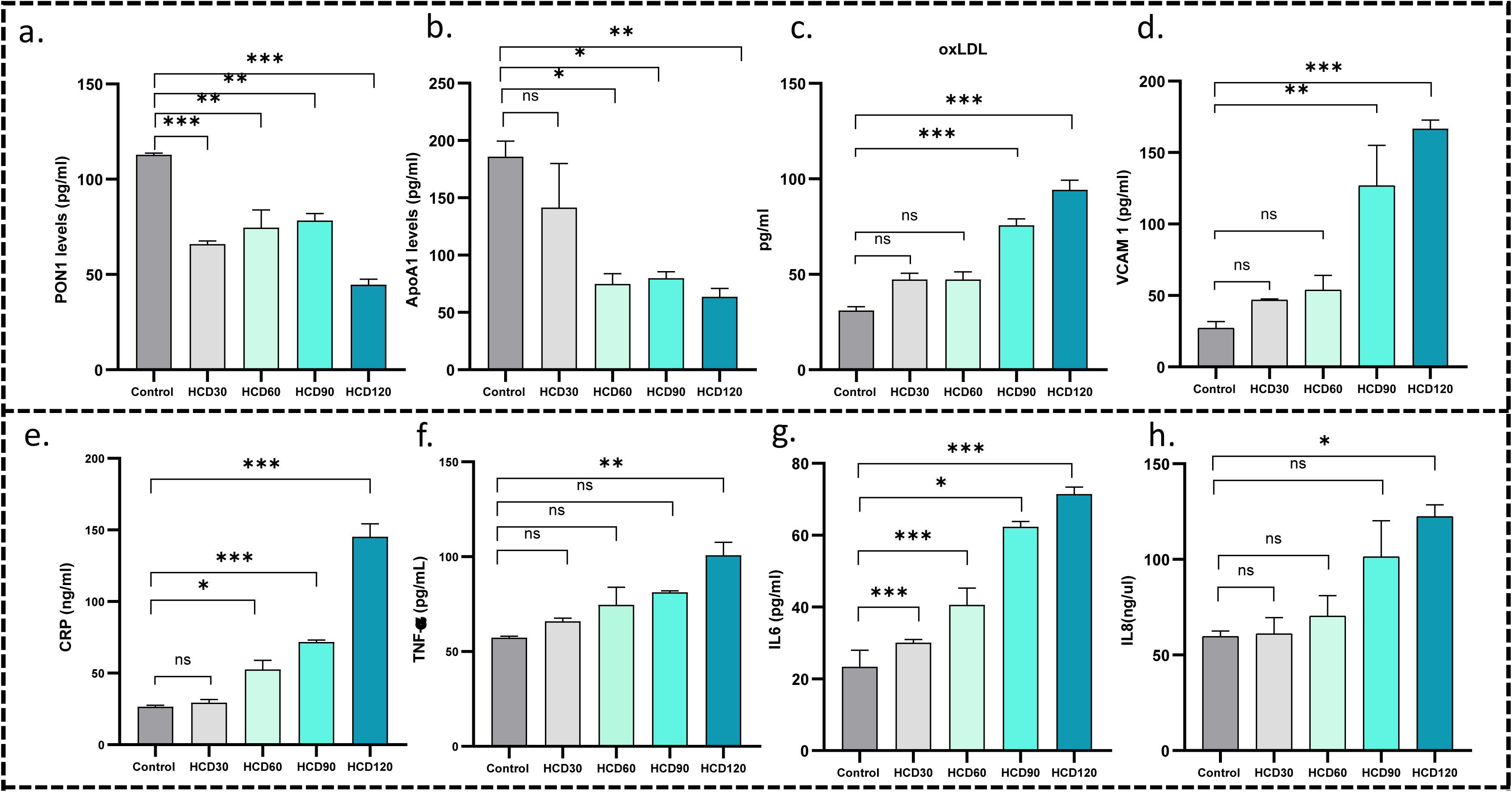
Levels of **a.** Paraoxonase 1, **b.** lipoprotein (Apolipoprotein A1) and **c.** oxidized LDL, **d.** Cell adhesion proteins (VCAM 1), inflammatory markers **(e.** CRP, **f.** TNF-α, **g.** IL-6, **h.** IL-8,) and in the serum of control and high cholesterol diet fed animals was performed using ELISA. Values are expressed as Mean ± S.E.M. for there or six independent experiments from each group. Statistical analysis was carried out using the one-way ANOVA followed by Tukey’s post-hoc test. (*p<0.5, **p<0.01, ***p<0.001)

**Figure 5.**
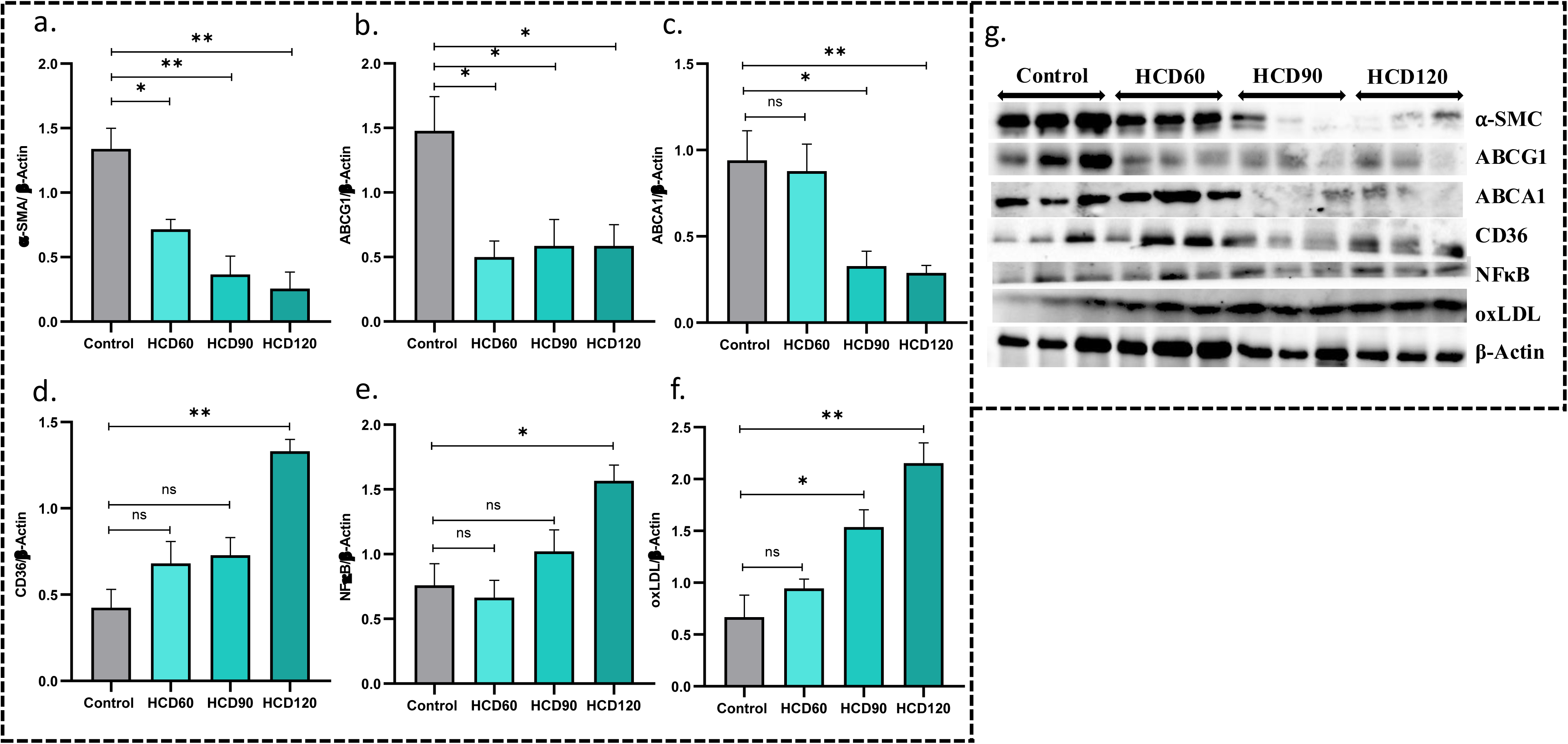
a-g. Western blotting results showing that the expression level of proteins and quantification of western blot band intensities by Image lab software (Bio-rad). Values are expressed as Mean ± S.E.M. for there or six independent experiments from each group. Statistical analysis was carried out using the one-way ANOVA followed by Tukey’s post-hoc test. (*p<0.5, **p<0.1)

**Figure 6.**
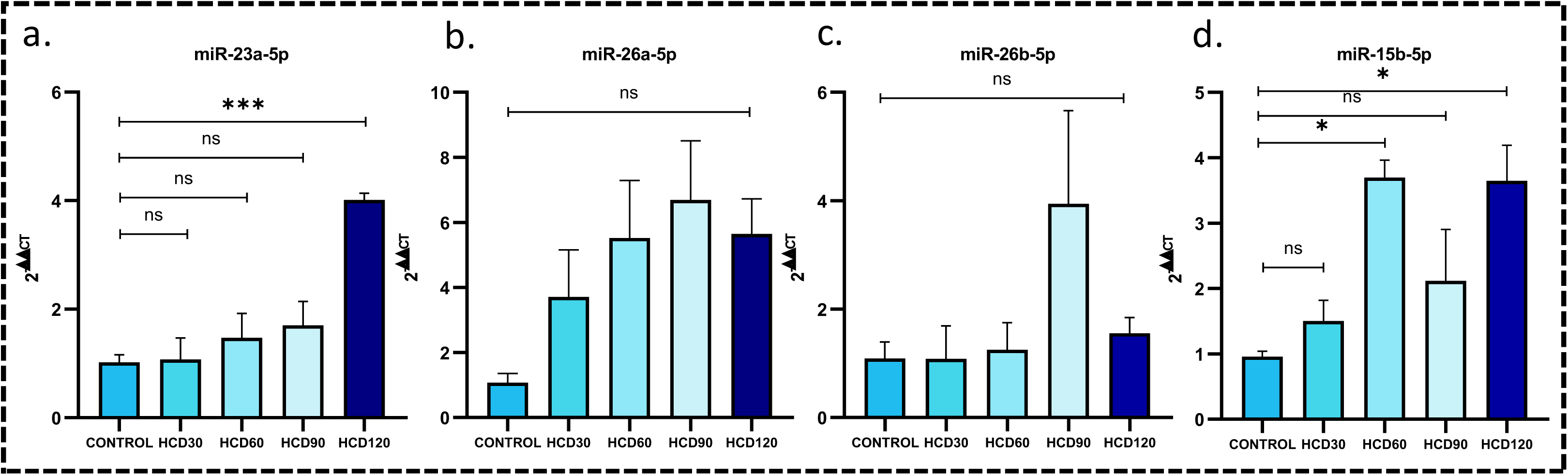
Serum miRNA Levels of High cholesterol diet (30days, 60days, 90days, 120days) fed rats. Graphs showing the relative fold change of miR-23a-5p, miR-26a-5p, miR-26b-5p and miR-15b-5p.

**Figure 7.**
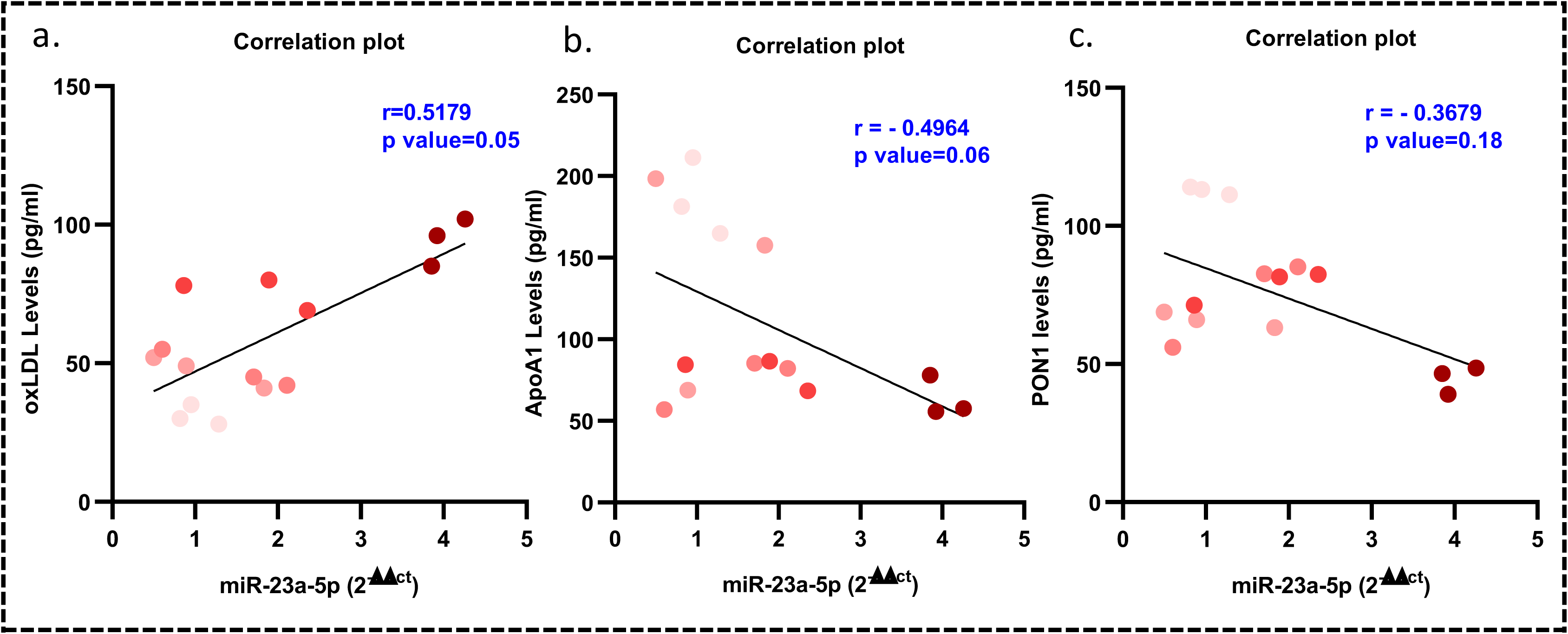
Spearman correlation between serum miR-23a-5p expression and oxidized LDL (oxLDL), ApoA1 and PON1 levels in rats subjected to varying durations of high cholesterol diet. A moderate positive correlation was observed (*r* = 0.5179, *p* = 0.05), indicating a possible link between oxidative lipid stress and miR-23a-5p regulation.

**Figure 8.**
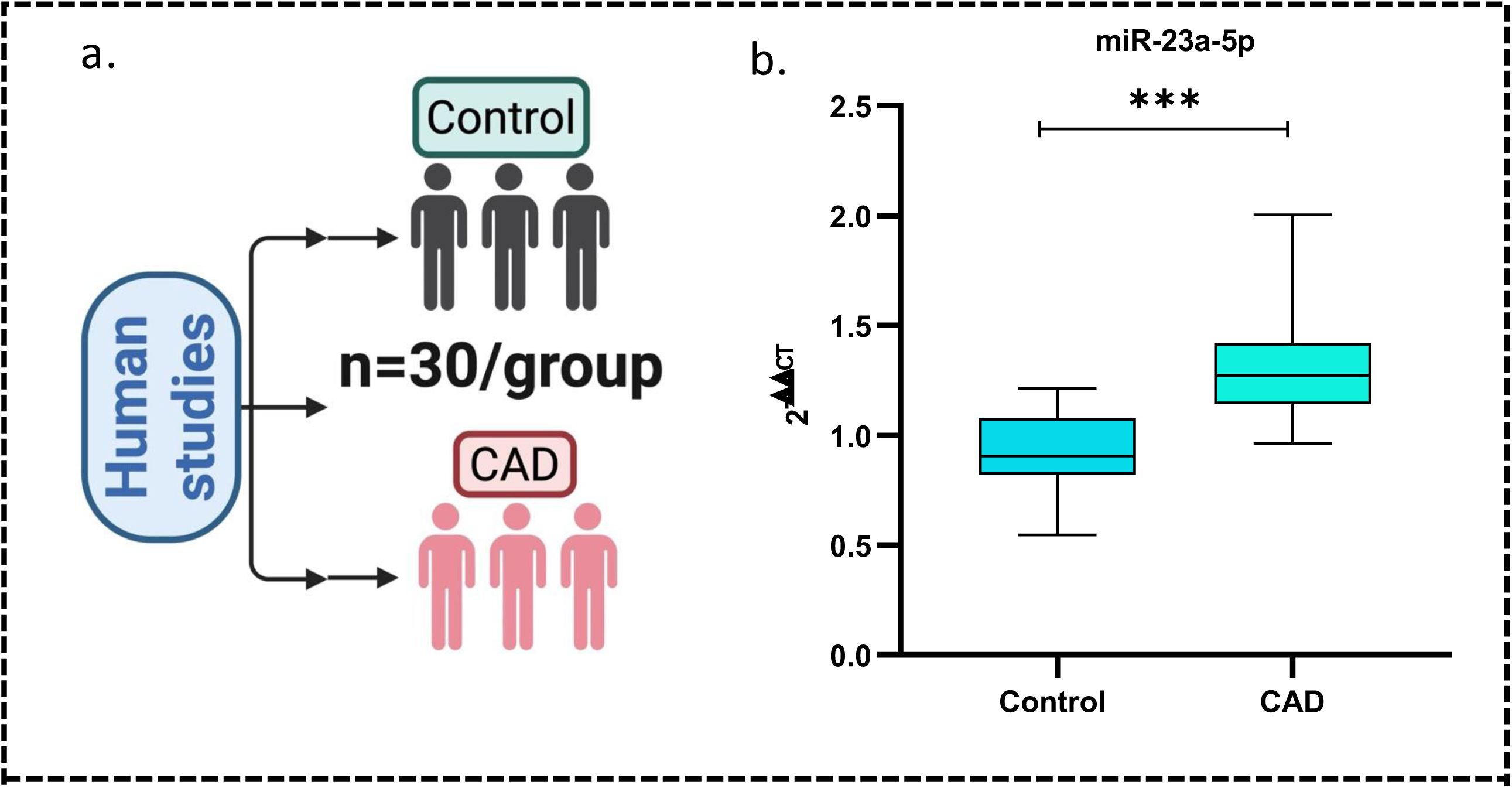
**a.** Number of human sample size used for the experiment. **b.** Serum miRNA Levels of healthy controls and CAD patients. Graph shows the relative fold change of miR-23a-5p

## Notes

### Competing Interest Statement

The authors have declared no competing interest.

